# The genetic diversity of Ethiopian barley genotypes in relation to their geographical origin

**DOI:** 10.1101/2021.11.10.468099

**Authors:** Surafel Shibru Teklemariam, Kefyalew Negisho Bayissa, Andrea Matros, Klaus Pillen, Frank Ordon, Gwendolin Wehner

## Abstract

Ethiopia is recognized as a center of diversity for barley, and its landraces are known for the distinct genetic features compared to other barley collections. The genetic diversity of Ethiopian barley likely results from the highly diverse topography, altitude, climate conditions, soil types, and farming systems. To get detailed information on the genetic diversity a panel of 260 accessions, comprising 239 landraces and 21 barley breeding lines, obtained from the Ethiopian biodiversity institute (EBI) and the national barley improvement program, respectively were studied for their genetic diversity using the 50k iSelect single nucleotide polymorphism (SNP) array. A total of 983 highly informative SNP markers were used for structure and diversity analysis. Three genetically distinct clusters were obtained from the structure analysis comprising 80, 71, and 109 accessions, respectively. Analysis of molecular variance (AMOVA) revealed the presence of higher genetic variation (89%) within the clusters than between the clusters (11%), with moderate genetic differentiation (PhiPT=0.11) and adequate gene flow (Nm=2.02). The Mantel test revealed that the genetic distance between accessions is poorly associated with their geographical distance. Despite the observed weak correlation between geographic distance and genetic differentiation, for some regions like Gonder, Jimma, Gamo-Gofa, Shewa, and Welo, more than 50% of the landraces derived from these regions are assigned to one of the three clusters.

## Introduction

Barley (*Hordeum vulgare* L.) ranks fifth in the acreage and production of cereals after tef, maize, wheat, and sorghum in Ethiopia. It accounts for 5.63% of the total cereal production (811,782.08 hectares (ha)) with a productivity of 2.18 ton/ha in 2018/19 [1]. It is a widely adapted crop, cultivated from drought prone lowlands of 1,500 meters above sea level to highlands of Ethiopia with an altitude of 3,400 meters above sea level with adequate moisture [2]. Most of the barley acreage is located in the altituted range of 2,400 to 3,400 meter above sea level in the northern and central part of the country [3]. In Ethiopia, barley is an important cereal crop grown by smallholder farmers for subsistence with limited capacity for modern agricultural practices, and in areas where soil fertility, drainage conditions, and topography are not suitable to produce other crops [4]. It is cultivated in two seasons; *‘meher’*, which is the major rainy season (June to October) in which diverse genotypes are grown, and *‘belg*’ with less amount of rain (late February to early July) in which most early maturing varieties are grown [5]. The total amount of barley production in *‘meher*’ is by far exceeding the one in *‘belg’*, which covered 84.5% of the total area of production and 93.0% of the total yearly barley harvest in 2013/14 [6].

Ethiopia is recognized as a center of diversity for barley, as it is cultivated in a wide range of agro-ecology zones for centuries, and its landraces have exhibited distinct genetic diversity from the rest of the world’s barley collections [7-9]. The presence of diversified and distinct genetic features have been explained by geographical isolation of the country from other barley growing regions for long periods together with the occurrence of diverse soil types, climate conditions, elevation, and landscape, which affect the type of farming system practices [10, 11]. One study indicated that Ethiopian barley population structure depends on the farming system, elevation, and barley row types [12]. Additionally, social factors like a preference of genotypes suited for different use also contributed significantly to the diversification [13]. Therefore, it was suggested that the diversity in Ethiopian barley landraces came due to a combination of long period accumulation of distant mutations, gene recombination, hybridization, natural selection, and human preference in a highly diversified agro-ecological environment [14].

The genetic resources of Ethiopian landraces are still rich and well maintained, as a report indicated that 95% of the Ethiopian smallholder farmers use landraces as the major seed source [15, 16]. Although barley is an inbreeding species with less than 5% of outcrossing, an increased rate of outcrossing was reported in Ethiopia, which is probably related to abiotic stress or variable environmental conditions [17]. Barley landraces at hand of farmers are genetically highly variable [18, 19], as farmers mainly focus to maintain morphologically uniform seeds than genetically uniform seeds, thus, sampling from smaller plots of farmers’ land may result in a collection of highly genetically diversified seeds [3].

Traditionally, farmers classified barley landraces based on kernel type as hulled, hull-less, and partially hulled barley [3]. Additionally, participatory research on durum wheat landraces revealed that farmers also considered yield, quality related to end-use products, and tolerance to different abiotic and biotic stresses like drought and diseases for the classification and selection of landraces [20]. Ethiopian barley landraces are particularly diverse in morphological appearance [21, 22] and bio-chemical composition, e.g. different hordein polypeptide patterns [23, 24] as well as anthocyanin coloration on seed coats, leaf sheath and stems [25].

The genetic structure of a population is influenced by variation in geographical collection distance, presence of geographical barriers like wetlands, mountains and gorges, as well as by the compatibility of genotypes to cross to each other. Besides this, the genetic structure is also due to the presence of barriers on the human local population over a long period of time [26].

Application of molecular tools improved the efficiency and precision of analysis of genetic relatedness in different crop species, as they helped to decipher whether the morphological, chemical, traditional and geographical classifications are in consistence to molecular structural analysis [27]. Different kinds of markers, i.e. AFLPs (amplified fragment length polymorphism), SSRs (simple sequence repeat), and SNPs (single nucleotide polymorphism) were used for genetic analysis of different cultivars, breeding lines and related species of barley [28-33]. Currently, SNP markers are commonly used to study genetic variation, as they are more abundant than other markers [34, 35]. The development of a 50k iSelect SNP array by [36] further enhanced the genetic exploration with accurate physical positions of the markers and detailed gene annotation.

The presence of genetic divergence between populations can be studied using Nei’s genetic distance [37]. Genetic abundance or richness within a population can be explored using the Shannon index [38, 39], whereas the variability within a population can be studied using heterozygosity indices [40]. The fixation index (FST) is widely used to investigate the genetic distance between populations [41, 42]. Using FST the gene flow between genetically distinct populations can be studied, while the gene flow between genotypes from different geographic locations can be studied using the Mantel test [43]. The neighbor-joining tree method is used to graphically demonstrate the distance between different genotypes based on their genetic background [44].

Several studies on the genetic distance of Ethiopian landraces using different molecular markers were conducted. Distinctive genetic features of Ethiopian landraces compared to other barley collections were reported, although a minimum genetic distance between different Ethiopian landraces was detected using RFPLs (restriction fragment length polymorphism) markers [45]. Another study revealed the presence of different levels of the allelic richness and genetic diversity in relation to altitude using seven SSR markers [19]. [46] also revealed a poor population structure for landraces collected from different regions of the country using 15 SSR markers. Genetic diversity studies of Ethiopian barley genotypes in relation to different world barley collections were also conducted using SSR [47], and AFLP markers [48] and the findings suggested Ethiopia as a second center of barley domestication.

Therefore, the aims of this study were, (i) to investigate the genetic diversity of Ethiopian barley landraces, and (ii) to analyses the role of the geographic origin, and defined agro-ecological zones in the formation of genetic structure using a highly informative 50k iSelect SNP array [36]. The outputs of the study will support the strategic collection and exploitation of existing barley genetic resources, to improve the livelihood of the subsistence farmers through strategic utilization of genetic resources available on the hand of smallholder farmers.

## Material and methods

### Plant material

A panel of 260 Ethiopian barley accessions was analyzed in this study (S1 Table). The 239 landrace accessions were obtained from the Ethiopian Biodiversity Institute. These were collected from diverse agro-ecological zones and represent different geographical regions of Ethiopia. The geographical locations in which the landraces were collected are shown in Fig 1, which is based on the GPS data of the collection area using the ArcGIS online web program (https://www.arcgis.com) [49]. Additionally, 21 barley breeding lines were obtained from the national barley improvement program of the Holetta Agricultural Research Center (HARC).

**Fig. 1.**
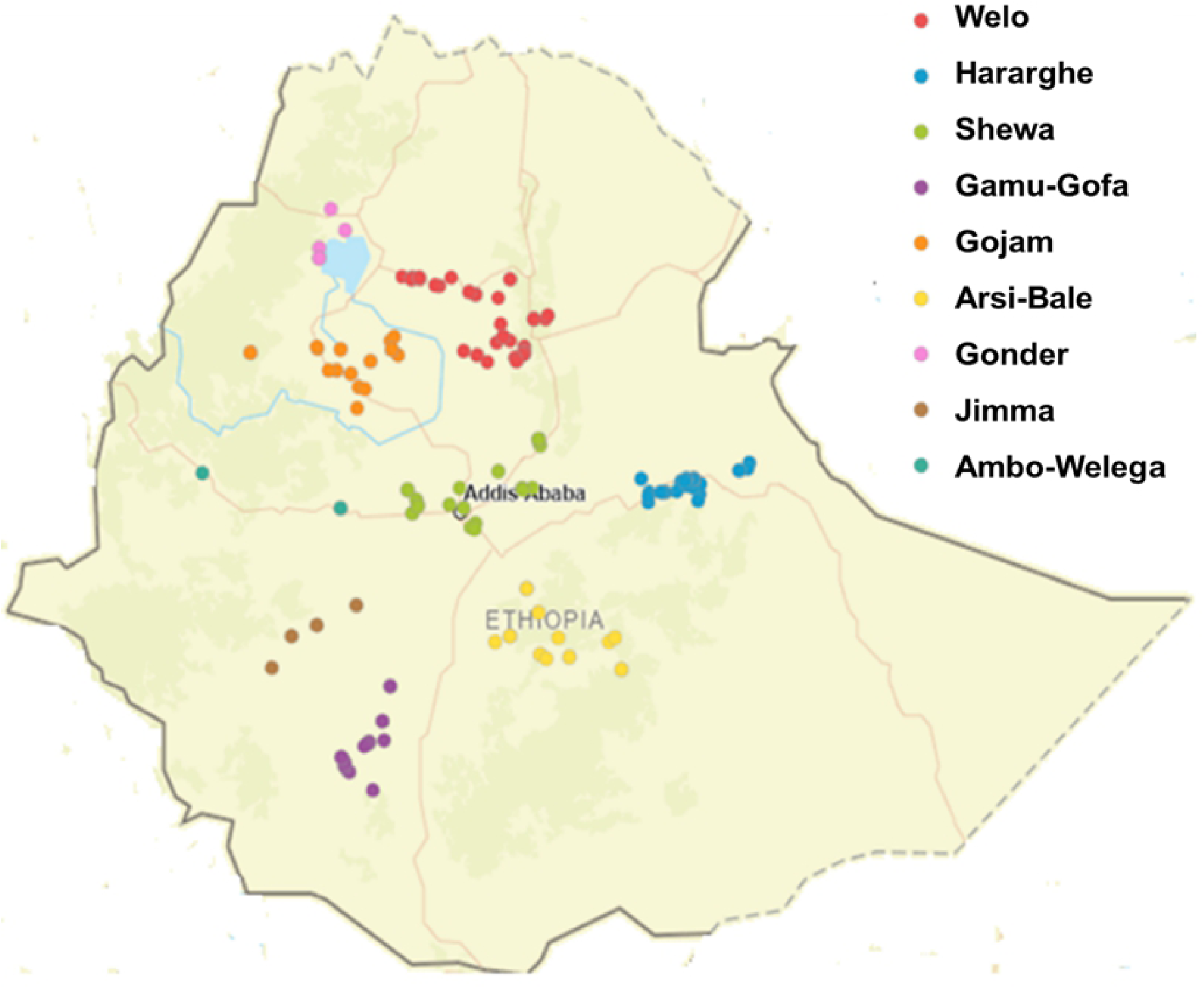
Ethiopian barley landrace accessions grouped by their geographical collection areas. Ethiopian boundary and geo-positions are indicated. Filled circles represent the 239 Ethiopian landraces collected at sometimes overlapping positions. Geographical positions are also detailed in S1 Table. The map was constructed using the online ArcGIS software suite vs. 10.8.1.

### Genotyping

Three seeds from each of the 260 accessions were grown in the greenhouse at day (16h)/ night (8h) temperatures of 20-22°C/17-19°C as described by [50] in multipot trays filled with Einheitserde ED73 soil containing 14% N, 16% P2O5 and 18% K2O in kg/m3 (H. Nitsch & Sohn GmbH & Co. KG, Germany). When plants had grown to the two to three leaf stage, leaf samples with an approximate size of 300 mg were taken from a single plant for genotyping. The genomic DNA was extracted using a modified CTAB (cetyltrimethylammonium bromide) method [51] and genotyped using the barley Illumina 50k iSelect SNP array [36] at TraitGenetics GmbH, Gatersleben, Germany.

An initial set of 40,387 markers was successfully extracted from genotyping. 10,644 SNP markers were obtained, after removing all monomorphic markers and imputation using Beagle [52] followed by final filtering using thresholds of 5% missing values, 3% minor allele frequency, and 12.5% heterozygous SNPs. A total of 983 highly informative markers were kept, using the software PLINK 1.9 (http://www.cog-genomics.org/plink/1.9/) [53], which uses the markers physical distances as well as pair wise linkage disequilibrium (LD) between adjacent markers to prune-in SNPs in strong LD, with unbiased representation along the genome.

### Population Structure

The 983 highly informative SNP markers were used for population structure and genetic diversity analysis. The population structure was calculated using the Structure software v.2.3.4 [54]. Computation of Bayesian statistical models was conducted by the Markov Chain Monte Carlo (MCMC) method based on 50,000 iterations following discard of 50,000 “burn-in” iterations. The web-based Structure Harvester software v0.6.94 (http://taylor0.biology.ucla.edu/structureHarvester/) [55] was used to identify the best probable number of subpopulation (k-value) according to [56]. From the best k-value, out of 10 replications the replication with the highest likelihood (mean LnP(K)) value was used as an inferred population cluster. The estimated membership coefficient of each accession was used to assign it to different clusters estimated by STRUCTURE based on the highest inferred cluster values. Principal coordinate analysis (PCoA) was applied to plot the population structure using the DARwin 5.0 software [57] based on the SNP matrix data.

### Genetic Diversity

The 983 highly informative SNP markers were used for genetic diversity analysis. AMOVA was performed based on the number of genetically distinct clusters obtained from the structure analysis. Information about genetic variation within and between clusters and gene flow (Nm) based on PhiPT (analogue of fixation index (FST)) were obtained from the analysis using the GenAlEX 6.5 software plugin for Excel [58]. The neighbor-joining tree, which is constructed based on the genetic distance of accessions [44], was created using the DARwin 5.0 software [57] to graphically demonstrate the presence of genetic distance between the subpopulations.

The genetic variance within and between clusters and gene flow (Nm) was calculated using the following formulas:

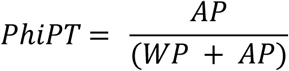

Where PhiPT is the genetic differentiation within and between clusters; AP is the estimated variance among clusters, and WP is the estimated variance within clusters.

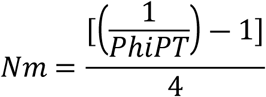

Where Nm is gene flow, and PhiPT is the genetic differentiation within and between clusters.

Genetic diversity indices i.e. Shannon’s information index (I), expected heterozygosity (He), unbiased expected heterozygosity (uHe), and percentage of polymorphic loci (PPL) were also calculated using frequency based analysis in the GenAlEX software [58]. Additionally, the Mantel test, which is used to estimate the gene flow by correlating the genetic distance with the spatial distance, i.e. GPS data in our case, was performed to get information on the genetic divergence across the geographical distance using the GenAlEX software [58].

## Results

### SNP analyses

From 43,461 scorable SNPs markers of the 50k iSelect SNP array [36]; 40,387 (92.9%) SNPs markers were successfully extracted in this experiment. However, 19,028 (47.1%) markers were immediately removed as monomorphic markers. From the remaining 21,355 markers, 10,767 SNPs markers (26.7% of the extracted set of markers) were removed by filtering for 3% minor allele frequency. Out of the 10,644 SNP markers, which were obtained after filtering, the highest number of markers was located on chromosome 2H (1857), and the least markers on chromosome 4H (1174). Similarly, for the 983 highly informative markers the highest number of markers was obtained for chromosome 2H (185), and the least for chromosome 4H (89) (Fig 2). The distribution of the markers revealed that most markers in the centromeric region were pruned-out, and the majority of the highly informative markers is located in the telomeric regions of all seven chromosomes (S1 Fig).

**Fig. 2.**
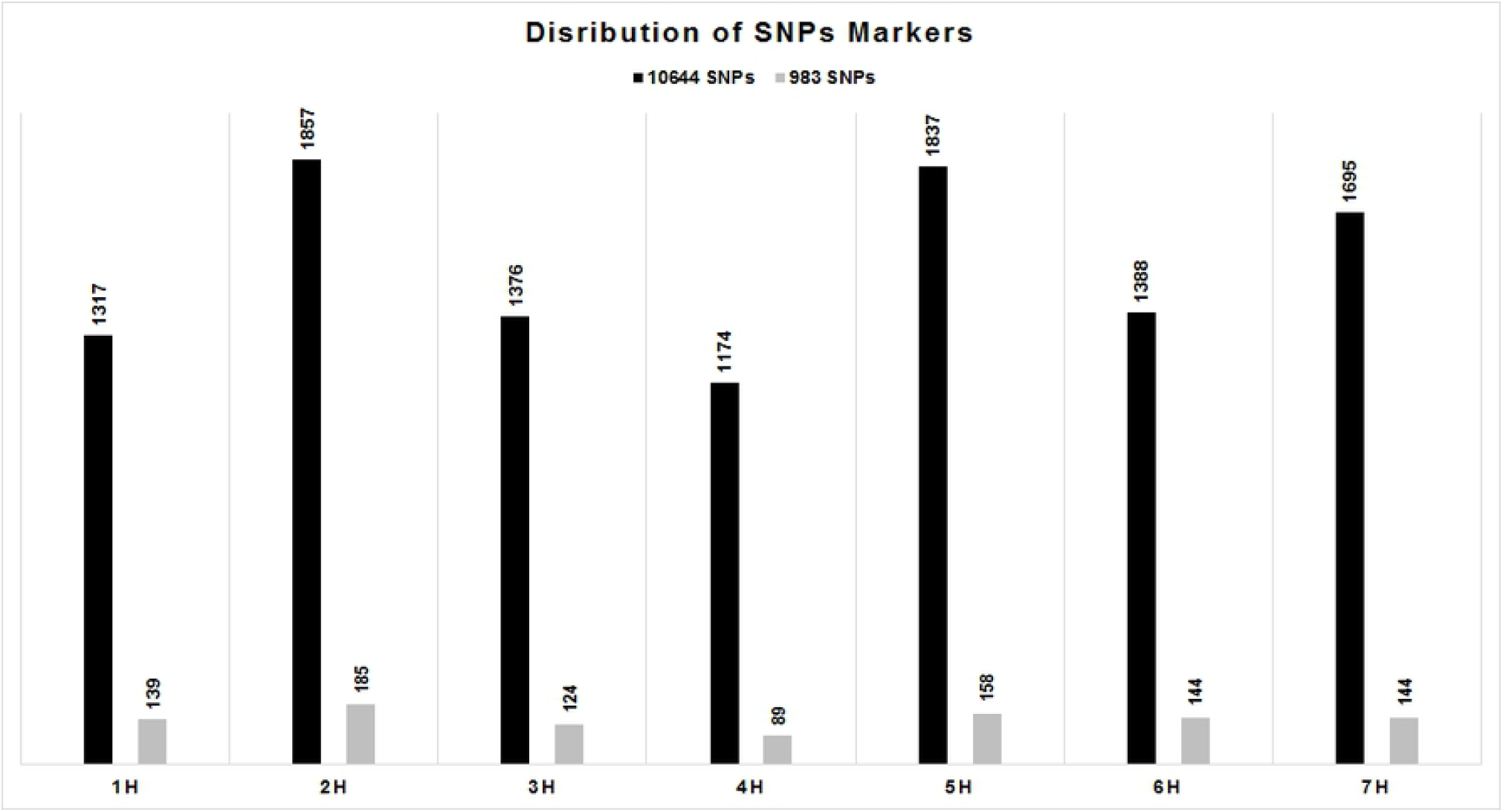
Distribution of filtered (10,644) and highly informative (983) SNPs across the barley chromosomes.

### Population structure analysis

Analysis of the population structure based on 983 SNP markers identified the best probable number of the subpopulation based on k-value at K=3, which therefore has been selected as an optimal number of inferred genetically defined clusters (Fig 3 a and b). According to the three genetically distinct clusters, cluster 1 consists of 80 accessions (30.8%), cluster 2 consists of 71 accessions (27.3%) and cluster 3 consists of 109 accessions (41.9%) out of the total of 260 accessions (Table 1). The average membership coefficient of the geographically defined populations indicated that Welo and Shewa population can be explained by cluster 1 and 2, respectively; whereas Gonder, Gamo-Gofa, and Jimma population were explained by cluster 3 (Table 2). When each member of a geographically defined population was re-assigned based on their highest probability value of the inferred clusters, 56% and 66% of Welo and Shewa accessions were clustered in genetically distinct cluster 1 and 2, respectively. Similarly, 88%, 86%, and 71% of Gonder, Gamo-Gofa, and Jimma accessions were grouped in the genetically distinct cluster 3, respectively (Table 1). Furthermore, 75% of the Ambo-Welega population was also assigned to cluster 3, but the low number of accessions has to be taken into account. Principal coordinate analysis (PCoA) indicated that PCoA1 and PCoA2 explained 5.87% and 4.88% of the variation, respectively. Despite these values being rather low, the high genetic variation within the set of accessions is reflected by the inferred three clusters (Fig 3c).

**Fig. 3.**
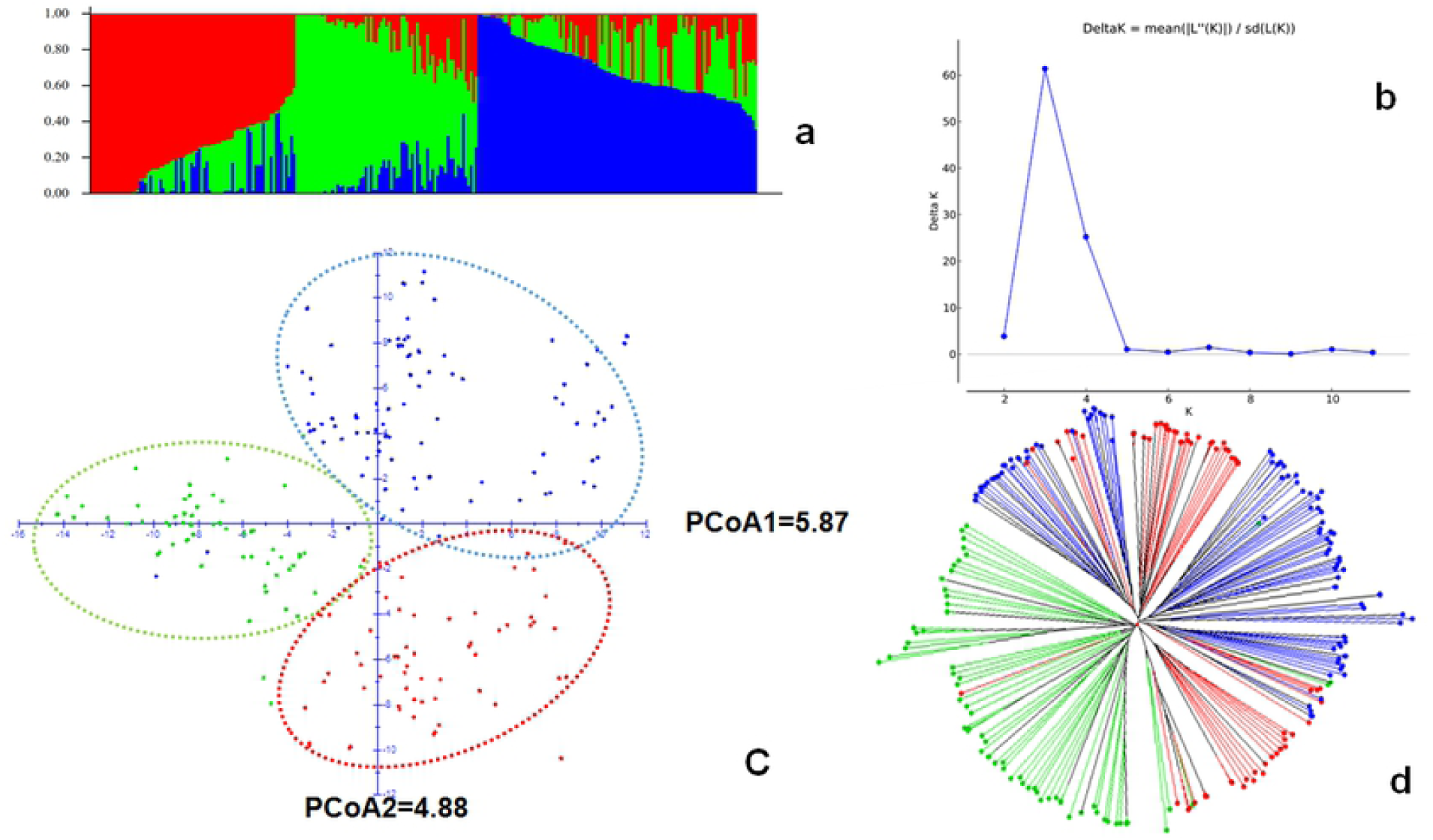
Population structure analysis for the 260 Ethiopian barley accessions. a) bar plot for estimated population structure of 260 Ethiopian barley accessions based on inferred three clusters (red = cluster 1, green = cluster 2, and blue = cluster 3); b) Structure harvester Delta K (ΔK) = 3; c) results of principal coordinate (PCoA) analysis, accessions were assigned based on their highest probability of inferred clusters; and d) weighted neighbor-joining tree for the structured subpopulations (red = cluster 1, green = cluster 2, and blue = cluster 3).

**Table 1.** Distribution of the Ethiopian barley accessions grouped by their geographical origin and based on the three genetically distinct clusters.

| Geographically defined subpopulations | Total accessions | Percentage of accessions in genetically distinct clusters |  |  |  |  |  |
| --- | --- | --- | --- | --- | --- | --- | --- |
|  |  | Cluster 1 |  | Cluster 2 |  | Cluster 3 |  |
|  |  | Number | % | Number | % | Number | % |
| Gonder | 8 | 1 | 12.5 | 0 | 0.0 | 7 | 87.5 |
| Arsi-Bale | 19 | 4 | 21.1 | 4 | 21.1 | 11 | 57.9 |
| Shewa | 38 | 6 | 15.8 | 25 | 65.8 | 7 | 18.4 |
| Ambo-Welega | 4 | 0 | 0.0 | 1 | 25.0 | 3 | 75.0 |
| Gojam | 28 | 6 | 21.4 | 14 | 50.0 | 8 | 28.6 |
| Welo | 59 | 33 | 55.9 | 13 | 22.0 | 13 | 22.0 |
| Gamo-Gofa | 28 | 2 | 7.1 | 2 | 7.1 | 24 | 85.7 |
| Jimma | 7 | 1 | 14.3 | 1 | 14.3 | 5 | 71.4 |
| Hararghe | 48 | 20 | 41.7 | 5 | 10.4 | 23 | 47.9 |
| HARC | 21 | 7 | 33.3 | 6 | 28.6 | 8 | 38.1 |
| **Total** | **260** | **80** | **30.8** | **71** | **27.3** | **109** | **41.9** |

**Table 2.** Average membership coefficient of Ethiopian geographically defined subpopulations based on the three genetically distinct clusters.

| Geographically defined subpopulations | Total accessions | Average membership coefficient of the subpopulations in the three genetically distinct clusters |  |  |
| --- | --- | --- | --- | --- |
|  |  | K1 | K2 | K3 |
| Gonder | 8 | 0.182 | 0.210 | 0.609 |
| Arsi-Bale | 19 | 0.202 | 0.355 | 0.444 |
| Shewa | 38 | 0.254 | 0.564 | 0.181 |
| Ambo-Welega | 4 | 0.221 | 0.421 | 0.359 |
| Gojam | 28 | 0.208 | 0.475 | 0.316 |
| Welo | 59 | 0.516 | 0.272 | 0.212 |
| Gamo-Gofa | 28 | 0.133 | 0.261 | 0.606 |
| Jimma | 7 | 0.145 | 0.256 | 0.600 |
| Hararghe | 48 | 0.437 | 0.161 | 0.402 |
| HARC | 21 | 0.334 | 0.376 | 0.290 |
| **Total** | **260** | **0.326** | **0.329** | **0.344** |

### Analysis of molecular variance (AMOVA)

AMOVA analysis was conducted based on the three genetically distinct clusters obtained through the analysis of population structure. The results revealed that variation within a cluster was accounting for higher variation (89%) than the variation among clusters (11%). The genetic differentiation was moderate (PhiPT = 0.11) with statistical significance at p < 0.001. Gene flow (effective migrant (Nm)) for the overall genetically distinct clusters was 2.02, which is characterized as a moderate rate of gene flow (Table 3).

**Table 3.** Analysis of molecular variance (AMOVA) for the Ethiopian barley accessions for the three genetically distinct clusters; PhiPT and Nm values for the total population

| Source | Degree of Freedom | Sum of square | Mean square | Estimated variance | Percentage of variation | $\Phi_{IPT}$ | $N_m$ |
| --- | --- | --- | --- | --- | --- | --- | --- |
| Among Populations | 2 | 6,576.5 | 3,288.2 | 35.3 | 11% | 0.11** | 2.02 |
| Within Populations | 257 | 73,143.0 | 284.6 | 284.6 | 89% |  |  |
| Total | 259 | 79719.5 |  | 319.9 | 100% |  |  |

\*\* p-value < 0.001

### Genetic diversity

The study of the genetic diversity indices of the three genetically distinct clusters indicate, that cluster 3 is more diverse than the other two clusters with values of I=0.47, He=0.31, uHe=0.31, PPL=99.1%, followed by cluster 2 (I=0.43, He=0.28, uHe=0.28, PPL=95.9%) while cluster 1 is the least divers one (I =0.39, He=0.26, uHe=0.26, PPL=88.2%) (Table 4).

**Table 4.** Genetic diversity indices for the genetically distinct clusters.

| Genetically distinct clusters | N | I | He | uHe | PPL |
| --- | --- | --- | --- | --- | --- |
| 1 | 80 | 0.39 | 0.26 | 0.26 | 88.2% |
| 2 | 71 | 0.43 | 0.28 | 0.28 | 95.9% |
| 3 | 109 | 0.47 | 0.31 | 0.31 | 99.1% |

“N” for number of observations, “I” for Shannon’s information index, “He” for expected heterozygosity, “uHe” for unbiased heterozygosity, and “PPL” for percentage of polymorphic loci

Generally, pairwise gene flow between the three genetically defined clusters is higher than one, which indicates the presence of adequate gene flow between these. Similarly, based on the results of pairwise PhiPT value, there is a moderate genetic differentiation between the subpopulations. The results indicate that the variation between genetically distinct cluster 1 and 2 is relatively larger (0.13) than between the other populations, whereas the gene flow between cluster 1 and 3 is higher (2.28) than between the other clusters (S3 Table).

The Mantel test, which is used to demonstrate the presence of spatial population structure indicated that the accessions were poorly structured, based on the GPS data of sampling with an R-squared value of 0.019 (Fig 4).

**Fig. 4.**
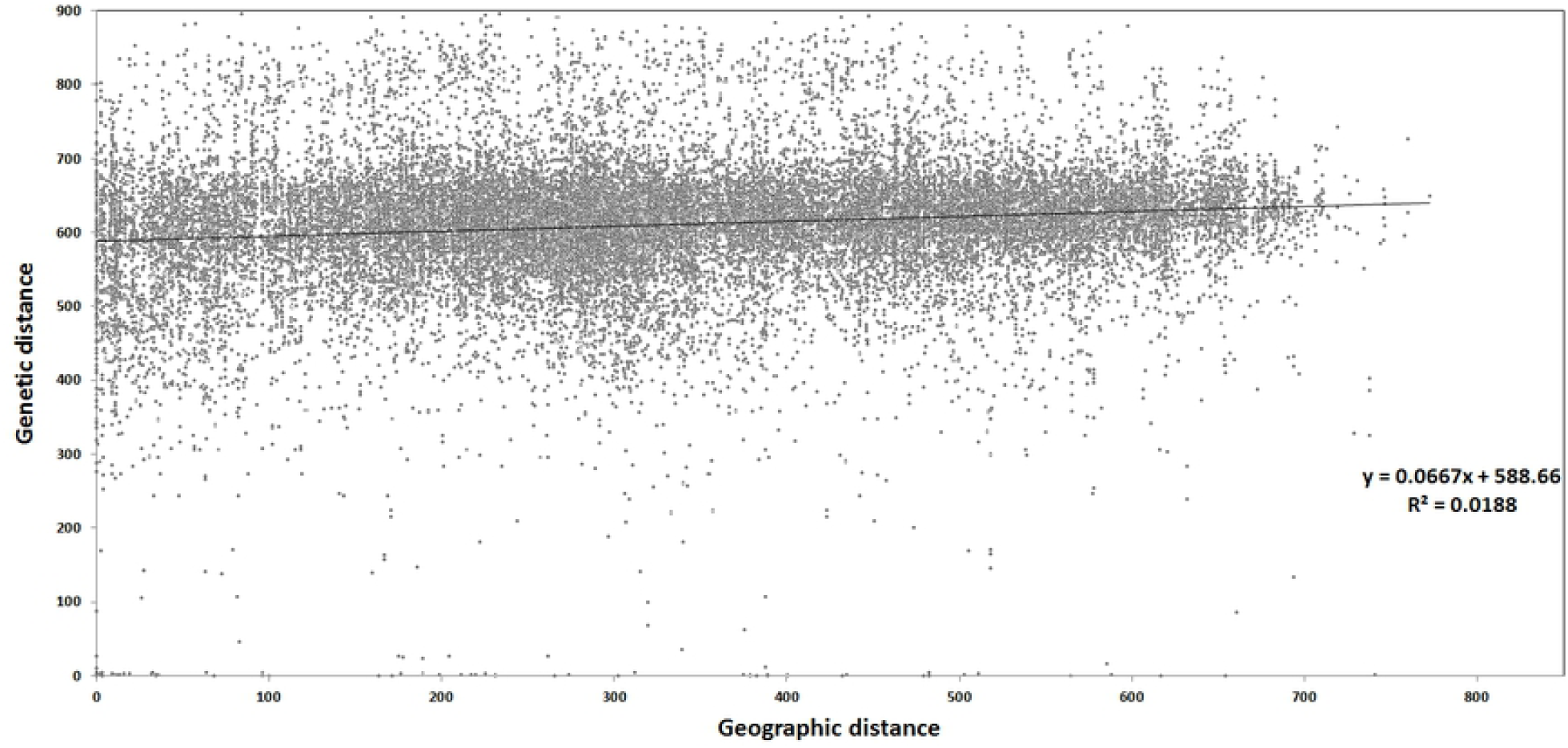
Mantel test for the 239 landraces based on the relationship between the genetic distance and the geographic distance based on GPS data.

## Discussion

From 43,461 scorable SNPs markers of the 50k iSelect SNP array [36]; the final number of SNPs markers (10,644) in our study was quite small compared to previous reports of 39,733 SNPs [59]; 33,818 SNPs [60] and 37,242 SNPs [61]. This may be explained by the under-representation of Ethiopian genotypes during the development of the 50k SNP array [36].

The distribution across the barley genome of the SNPs markers obtained after filtering was compared with the one of the 50k SNP array [36]. The genome regions containing the first and the second highest number of SNP markers were on chromosome 5H (8123) and chromosome 2H (7227) for the 50k SNP array development, whereas in our study the first and second highest representation were recorded on chromosome 2H (1857) and chromosome 5H (1837). The two genome regions with the least number of SNP markers were chromosome 1H (4828) and chromosome 6H (5441) for the 50k SNP array, while in this study chromosome 4H (1174) and chromosome 1H (1317) were least represented. Therefore, we considered the distribution of SNP markers along the seven barley chromosomes as similar with the 50k SNP array. A total of 983 highly informative markers, located in the telomeric regions of all seven chromosomes (S1 Fig), were kept for the population structure analyses.

According to the average membership coefficient, the predefined Welo and Shewa subpopulations were classified as genetically distinct in cluster 1 and 2, respectively (Table 1). By the ratio of accessions assigned in each cluster, accessions from Gonder, Gamo-Gofa and Jimma, predefined as subpopulations, appeared to be represented by cluster 3 (Table 2). Similarly, the average membership coefficient of the Gonder, Gamo-Gofa, and Jimma (Table 1) populations clearly suggested that they are members of cluster 3. [62] reported that landraces obtained from Shewa, Gonder, and Gojam have had minimum admixture, whereas landraces obtained from Arsi-Bale, Harerghe, and Welo were showing the highest ratio of admixture. Accordingly, in our study, landraces from Gonder, and Shewa were grouped in cluster 1 and 3 respectively; and Arsi-Bale and Harerghe were not defined by any cluster (Table 2).

Estimation of the population structure along the geographical and agro-ecological arrangement gives an important view on the pattern of population structure. In Ethiopia, studies conducted on different cereal crops highlighted the presence of higher genetic variation within geographical locations and altitude ranges for barley [46, 63, 64], durum wheat [65-67], and sorghum [68]. Similarly, the presence of minimum geographical structure was observed using the Mantel test in this study (Fig 4). This may be due to the fact that accessions from distantly located regions, i.e. Gonder, Jimma and Gamo-Gofa are grouped in cluster 3. Further analysis of AMOVA based on the agro-ecological zones of the accessions as a predefined subpopulation provided only 3% variation between agro-ecologies (S2 Table), although the variation between genetically distinct clusters was 11% (Table 3). On the contrary, previous genetic diversity studies on Ethiopian barley landraces suggested that the landraces’ population structure is dependent on the altitudinal gradient; which is mainly used for the classification of Ethiopian agro-ecologies [12, 19], but of which a minimum of variation explained was found in the current study (S2 Table). Although [62] reported the presence of weak association of structured populations of Ethiopian barley landraces with their geographical distance and climate conditions; in our study geographic locations slightly contributed to the variation among the structured populations in contrast to the agro-ecological conditions.

An overall estimation of gene flow for the three genetically distinct clusters was 2.02, which is greater than 1, and thus indicates the presence of gene flow between the subpopulations [26, 69, 70]. The overall population genetic differentiation (PhiPT) value is (0.11) indicating the presence of moderate differentiation between the genetically clustered subpopulations [41]. Similarly, the pairwise PhiPT value between clusters ranges from 0.10 between cluster 1 and 3 to 0.13 between cluster 1 and 2 (S3 Table).

The presence of gene flow between the different genetically distinct clusters hints to the exchange of adaptive traits among them [26, 71]. The presence of weak geographical or agro-ecological structure for Ethiopian barley landraces [46, 63, 64] may be explained by the exchange of important adaptive genetic traits between different genetically distinct clusters. The 21 breeding lines used in the study are proportional distributed in the three clusters (Table 1 and Table 2), which is also an indicator, that the national breeding program is introducing important adaptive traits from landraces in new varieties. In our study, an adequate gene flow between different clusters was observed (S3 Table). Similarly, gene flow > 1 (Nm=2.95), was reported by [63], using Ethiopian barley landraces collected from different regions. From these results, we concluded that the presence of an adequate amount of gene flow among different subpopulations contributes to a wider adaptation of Ethiopian landraces.

From the three genetically distinct clusters, cluster 1 is explained by the Welo predefined subpopulation. [72] described that around the Welo location barley is an important crop, and farmers conserve the landraces for different reasons, such as for their suitability to use it for short and long rainy seasons (maturity), yield potential, tolerance to water logging, frost and low soil fertility, social preference (taste and visual appearance), and storability. Furthermore, barley is also used as a main dish (to prepare *injera*, and bread) in this area, and a special dish and beverage (*tihlo* and *korefe*), which are exclusively prepared from barley, are commonly consumed in this area [72]. Thus, another assumption for the formation of this genetically clustered population may be related to the landraces quality to prepare staple food as well as special dishes and beverages.

Cluster 3 mainly contains landraces from Gamo-Gofa, and the production of barley in Gamo-Gofa is mainly on highlands with an altitude higher than 2,500 meter above sea level [73]. Such highland topographies are characterized by having low road access to connect with nearest commercial cities. As a result the diversity in such areas will be kept unchanged. Accordingly, studies suggested an increased market access in the community contributing to an increase in crop diversity [74, 75]. In our study, the presence of low market access likely contributed to the grouping of 86% of Gamo-Gofa accessions in cluster 3. Although farmers varieties selection criteria in Gamo-Gofa are similar to other locations, barley is not served as main dish in the region and usually used to prepare special dishes and beverage (local beer) during a festive holiday and special occasions [76]. We therefore assume that the farmers selection criteria for varieties may be based on the end use of the product, and consequently landraces in cluster 3 might be related with such quality traits.

Shewa is located in the central part of Ethiopia, with best road facilities, and high consumer demand. Farmers usually produce barley for home consumption and market; and [77] reported that farmers produce barley as it is adapted very well comparing to other cereal crops to the low fertility soil in this region. Barley is used in this region to prepare local liquor and local beers, which have great demand for market. Additionally farmers produce suitable landraces to prepare the main dish (*injera*) [77]. A significant reduction in the number of farmer’s varieties comparing to the previous time was reported in Shewa [78] due to socio-economic and environment related reasons. Such genetic erosion may not just be a recent history in the region, but might also be present in the previous decades, which is ultimately narrowing the genetic bases of the landraces in this area. The result obtained from weighted neighbor-joining tree (Fig 3d) and the pairwise gene flow (Nm) and PhiPT (S3 Table) indicated that cluster 2 derived from slightly different predecessor families, in comparison to cluster 1 and 3 which are closer related. Therefore, the remaining landraces around Shewa with a narrow genetic base may be mostly related to cluster 2 (Fig 3d, Table 2).

Cluster 3 is a diverse cluster based on the results of genetic indices (Table 4). 86% of accessions from Gamo-Gofa are assigned to this cluster and [79] also described that landraces obtained from Gamo-Gofa region have higher diversity index compared to other regions. On the contrary, landraces from Gonder, which are also grouped in cluster 3, have been described for having least diversity in that study.

## Conclusions

Genetic structure and diversity of 260 Ethiopian barley landraces, comprising 239 accessions from Ethiopian Biodiversity Institute, and 21 barley breeding lines of the national barley improvement program of the Holetta Agricultural Research Center, were investigated based on data obtained from the barley 50k iSelect SNP array. The presence of higher rates of monomorphic markers with minor allele frequency less than three seems characteristic for Ethiopian barley accessions compared with other barley collections from the world. AMOVA revealed the existence of high genetic diversity within genetically distinct populations in comparison to the genetic diversity between genetically distinct populations. This may be due to the minimum geographical structure of landraces and the presence of higher gene flow between accessions originated from distant geographic locations. The use of barley for different food recipes and beverages may also play a role in the genetically clustered population structure as [13] described the use of different barley types for different purposes by the society of different regions. However, further analysis based on the nutritional quality of each landraces in specific geographical locations may be required. Our results will support the strategic collection and exploitation of the existing genetic resources of Ethiopian barley landraces, and will help improving farm management of subsistence farmers through the dedicated utilization of genetic resources in the near future.

## Acknowledgments

The authors would like to acknowledge the Federal Ministry of Food and Agriculture (BMEL, Bundesministerium für Ernährung und Landwirtschaft), Germany for funding of this research project (FKZ 2813FS01). The Ethiopian Biodiversity Institute (EBI) and the Holetta Agricultural Research Center (HARC) are acknowledged for providing the initial barley study materials. We also like to thank the laboratory and greenhouse technicians and the administrative officers of the Julius Kühn Institute (JKI) for their technical and administrative facilitation of the research work. We acknowledge Heike Lehnert for her assistance and guidance in genomic data analysis.

## Author contribution statement

The research idea was developed by FO and GW. All authors contributed to the study conception and design. Material preparation, data collection, and analyses were performed by SST, KNB, and GW. Supervision was done by FO, KP, AM and GW. Data analysis and the first draft of the manuscript was written by SST, and all authors commented on previous versions of the manuscript. All authors read and approved the final manuscript.

## Supporting information

**S1 Figure. Physical map distribution of SNP markers across the seven barley Chromosomes. A: Filtered 10**,**644 SNP markers; B: Highly informative 983 SNP markers**.

**S1 Table. Geographical location and agro-ecological zones of the Ethiopian barley landrace collection**.

**S2 Table. Molecular variance (AMOVA) for the Ethiopian barley accessions based on the 14 defined agro-ecological zones; genetic differentiation (PhiPT) and gene flow (Nm) values of the total population**.

**S3 Table. Pairwise correlation matrix for genetic differentiation (PhiPT) and gene flow (Nm) values between the three genetically distinct clusters, whereas shaded cells represent Nm values, unshaded ones represent PhiPT values**.

